# Discovery of the world’s highest-dwelling mammal

**DOI:** 10.1101/2020.03.13.989822

**Authors:** Jay F. Storz, Marcial Quiroga-Carmona, Juan C. Opazo, Thomas Bowen, Matthew Farson, Scott J. Steppan, Guillermo D’Elía

## Abstract

Environmental limits of animal life are invariably revised upwards when the animals themselves are investigated in their natural habitats. Here we report results of a scientific mountaineering expedition to survey the high-altitude rodent fauna of Volcán Llullaillaco in the Puna de Atacama of northern Chile, an effort motivated by video documentation of mice (genus *Phyllotis*) at a record altitude of 6205 m. Among numerous trapping records at altitudes >5000 m, we captured a specimen of the yellow-rumped leaf-eared mouse (*Phyllotis xanthopygus rupestris*) on the very summit of Llullaillaco at 6739 m. This summit specimen represents an altitudinal world record for mammals, far surpassing all specimen-based records from the Himalayas and elsewhere in the Andes. This discovery suggests that we may have generally underestimated the altitudinal range limits and physiological tolerances of small mammals simply because the world’s highest summits remain relatively unexplored by biologists.

## Introduction

The environmental limits of animal life have always fascinated biologists, and new discoveries about organismal adaptability continually force us to revise our assumptions about such limits. At high altitude, endothermic vertebrates are forced to cope with a combination of environmental stressors, the most salient of which are the reduced partial pressure of oxygen (hypoxia) and freezing temperatures. Nonetheless, numerous alpine mammals and birds have evolved physiological capacities for meeting such challenges^1–6^ and are capable of surviving at surprisingly lofty altitudes so long as food is available.

Upper altitudinal limits of wild mammals in the Himalayas and Andes are generally thought to fall in the range 5200-5800 m above sea level^7–12^. Such limits are surely dictated by food availability in addition to physiological capacities for tolerating hypoxia and extreme cold. The altitudinal range limits of alpine birds and mammals are often not known with certainty due to scanty survey data in inaccessible highland regions, and many published records in the scientific literature are surpassed by sightings reported by members of mountaineering expeditions. For example, members of an Andean mountaineering expedition reported sightings of mice at >6500 m on Volcán Llullaillaco (Christian Vitry, pers. comm.), an altitude that far exceeds all specimen-based records in the region^7,11,13,14^. Llullaillaco (6739 m) is the second-highest active volcano in the world and straddles the border between Chile and Argentina.

Motivated by reported sightings of mice living at record altitudes, we organized a scientific mountaineering expedition to survey the rodent fauna of Volcán Llullaillaco and the surrounding altiplano/Puna de Atacama of northern Chile. Our world-record trapping results challenge current thinking about physiological and ecological constraints on the altitudinal range limits of mammals and indicate that the world’s highest summits are not as barren as once believed.

## Results and Discussion

On a 2013 mountaineering expedition to Volcán Llullaillaco, we filmed a mouse (identified as *Phyllotis* spp.) scurrying across a snowfield at 6205 m above sea level (24°43.052’S, 68°33.323’W)(Supplementary Video 1), an altitude that far surpasses existing records for wild mammals. This sighting motivated our subsequent high-altitude trapping expedition in February 2020. During the course of this expedition, we live-trapped rodents from ecologically diverse sites on the altiplano and puna spanning >4300 m of vertical relief (Fig. 1, Supplementary Table 1). On Volcán Llullaillaco (Fig. 2), we live-trapped rodents in and around Aguadas de Zorritas (4140-4360 m), base camp – Ruta Normal (4620 m), base camp – Ruta Sur (5070 m), high camp – Ruta Sur (5850 m), and the volcano summit (6739 m). In total, we collected museum voucher specimens of 80 mice representing four species: Andean altiplano mouse (*Abrothrix andina*), altiplano laucha (*Eligmodontia puerulus*), yellow-rumped leaf-eared mouse (*Phyllotis xanthopygus*), and Lima leaf-eared mouse (*P. limatus*). We collected *Eligmodontia puerulus* and *Abrothrix andina* at maximum altitudes of 4099 and 4620 m, respectively; these altitudes approximate or exceed previous records for these species^15–16^. Our altitudinal records for *Phyllotis limatus* and *P. xanthopygus* (5070 and 6739 m, respectively) far exceed existing records for both species^11,13–14,17–18^.

**Figure 1.**
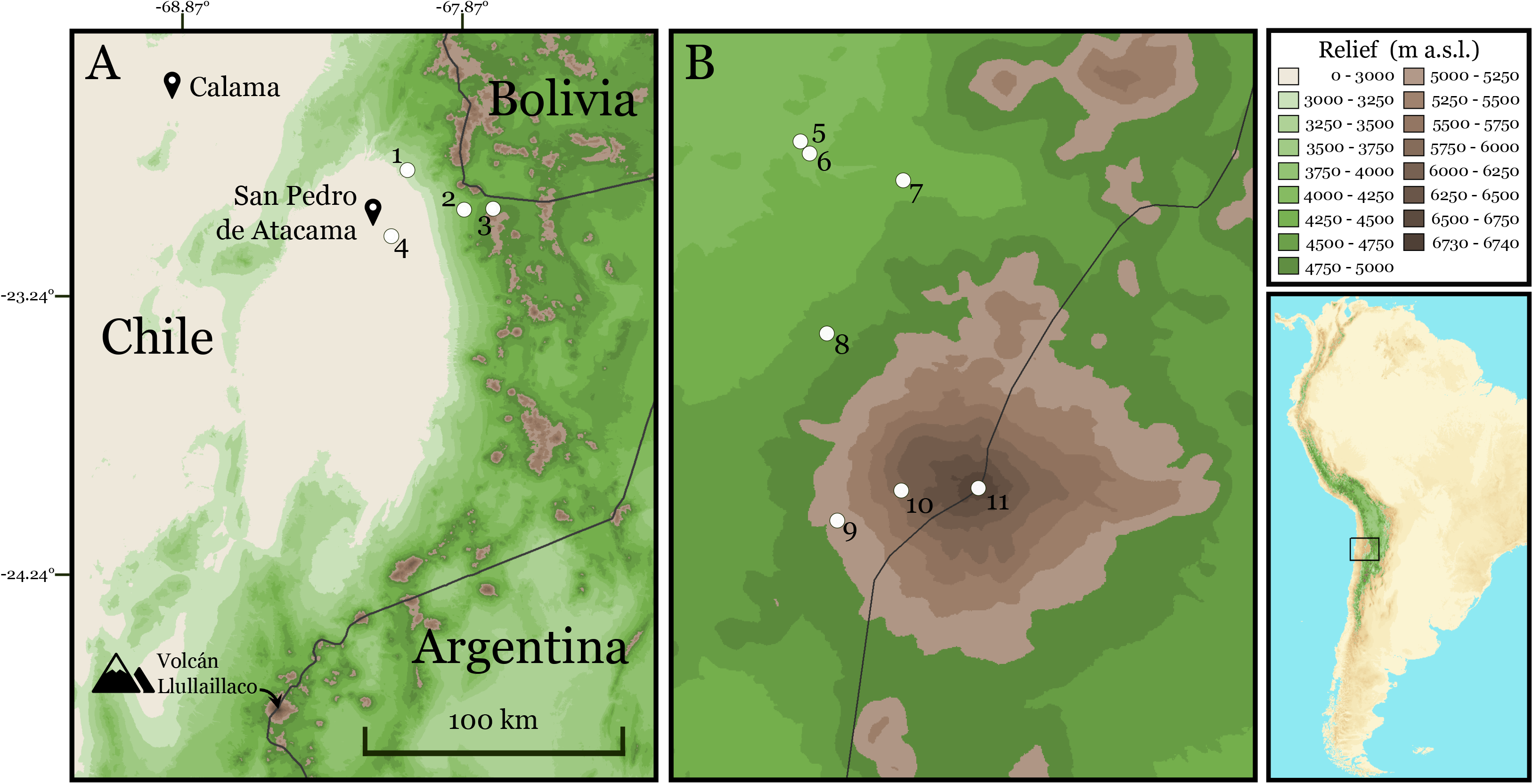
Map showing collecting localities in the altiplano and Puna de Atamaca, including Volcán Llullaillaco, Región de Antofagasta, Chile.

**Figure 2.**
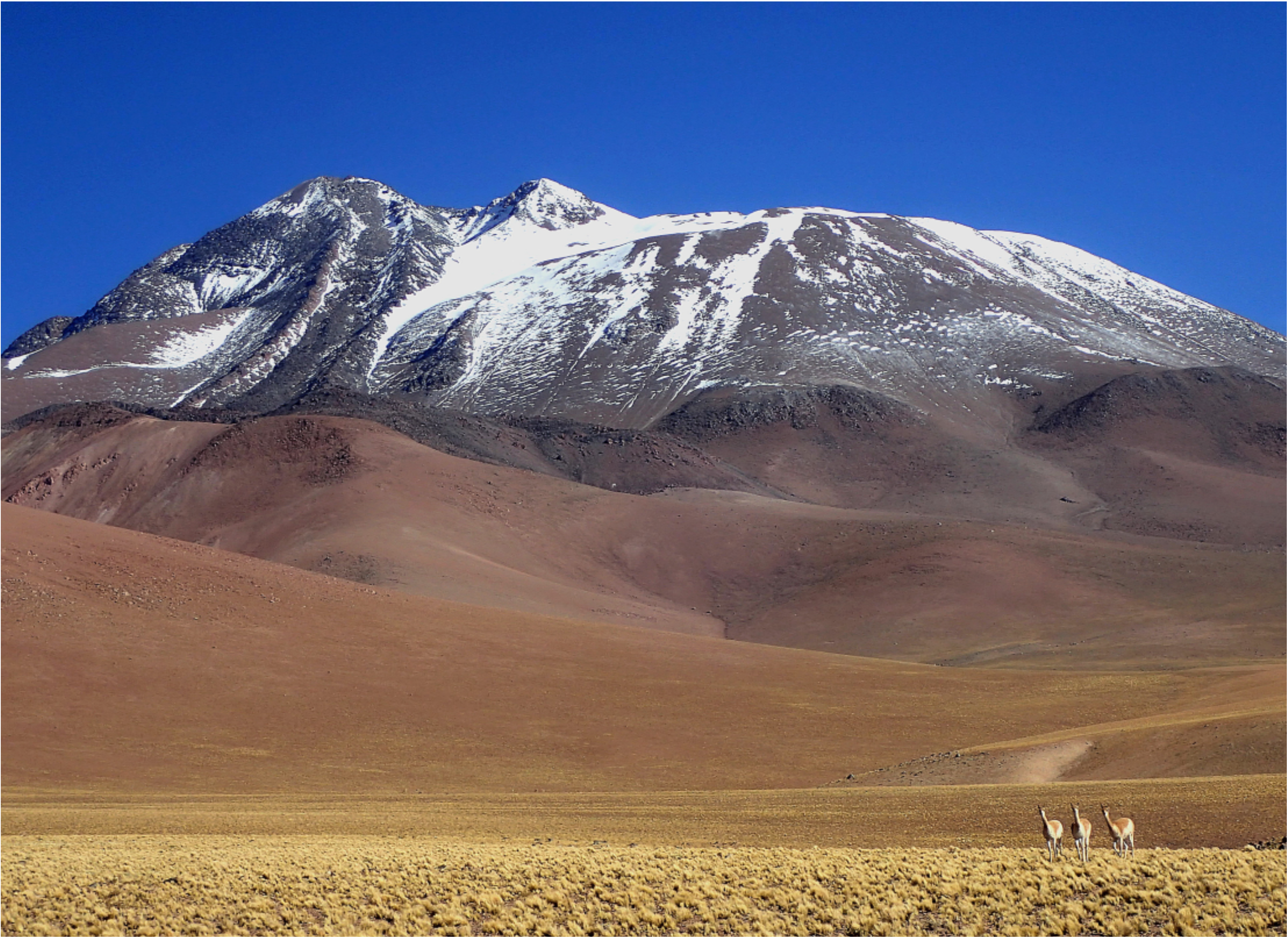
View of Volcán Llullaillaco (6739 m) from the west, Región de Antofagasta, Chile.

We captured the 6739 m specimen of *Phyllotis xanthopygus* on the very summit of Llullaillaco (Figs. 3, Supplementary Video 2). This summit specimen (Fig. 4) represents an altitudinal world record for mammals, far surpassing all specimen-based records from the Himalayas and elsewhere in the Andes.

**Figure 3.**
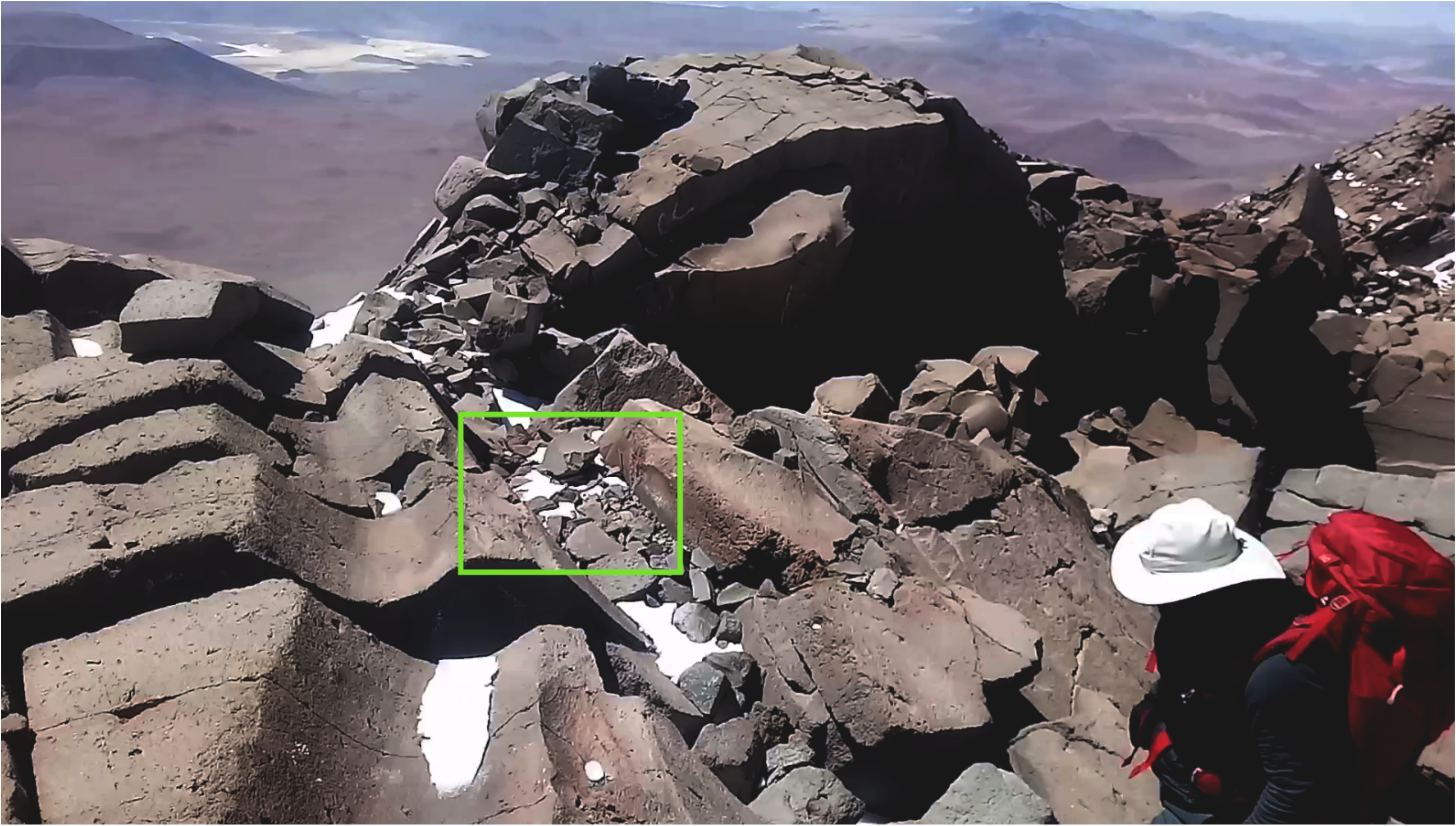
Site of capture of the record specimen of *Phyllotis xanthopygus* (GD2097) on the summit of Volcán Llullaillaco (6739 m), Región de Antofagasta, Chile (24°43.235’S, 68°32.208’W).

**Figure 4.**
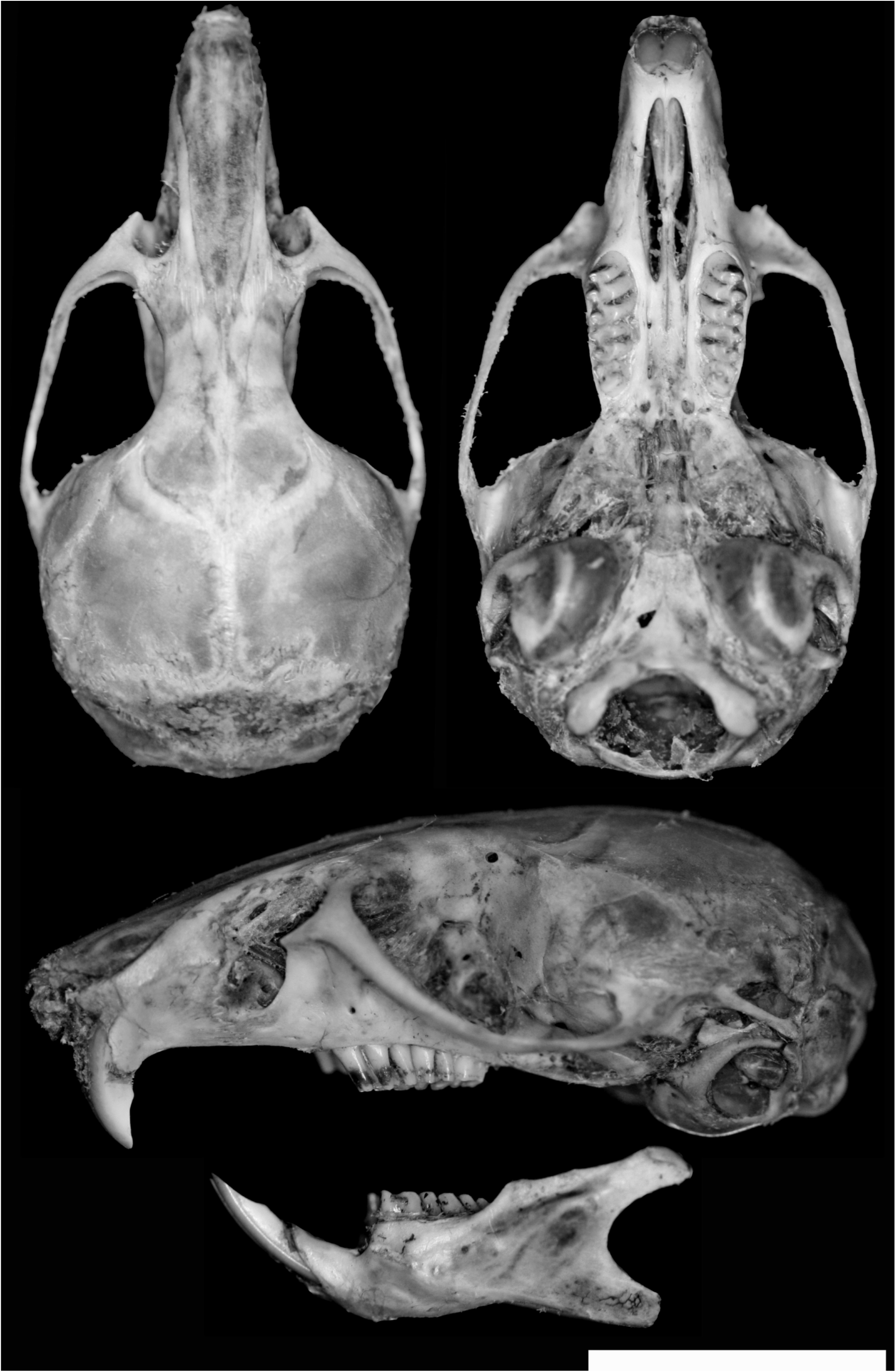
Dorsal, ventral, and lateral views of skull and lateral view of mandible of *Phyllotis xanthopygus rupestris* (adult male, Universidad Austral de Chile, GD2097) collected 18 February 2020 from the summit of Volcán Llullaillaco (6739 m), Región de Antofagasta, Chile. Scale bar = 10 mm.

Phylogenetic analysis of *cytochrome b (cytb*) sequences confirmed the species identifications of our record specimens of *Phyllotis limatus* and *P. xanthopygus* and revealed close relationships with conspecific specimens that were previously collected in the altiplano and adjacent regions of northern Chile and Argentina (Fig. 5). The *cytb* haplotype of our summit specimen of *P. xanthopygus* (GD 2097) groups with those of previously collected altiplano specimens of *P. xanthopygus rupestris*^19^. Moreover, the *cytb* haplotype of this summit specimen is identical to that of another *P. x. rupestris* specimen (LCM1780) collected at Toconao, Chile, a locality ca. 180 km NNE of Volcán Llullaillaco. Similarly, two other specimens of *P. x. ruprestris* collected at different altitudes (GD 2082 at 4406 m and GD 2095 at 5069 m) on Volcán Lullaillaco share identical *cytb* haplotypes with a specimen (LCM 1737) collected at the mouth of the Loa River on the Pacific coast, ca. 400 km NW of Volcán Lullaillaco. Thus, not only does *P. x. ruprestris* range from sea level to the crest of the Andean Cordillera at 6739 m (the broadest altitudinal distribution of any mammal), but individuals with identical *cytb* haplotypes are found at opposite ends of this vast range. Three other sequenced specimens of *P. x. rupestris* from Volcán Llullaillaco (GD 2083, GD 2084, GD 2093) possess unique *cytb* haplotypes. Future studies based on a denser taxon sampling and additional loci will permit more detailed inferences regarding the demographic history of high-altitude *P. x. rupestris* on Volcán Llullaillaco.

**Figure 5.**
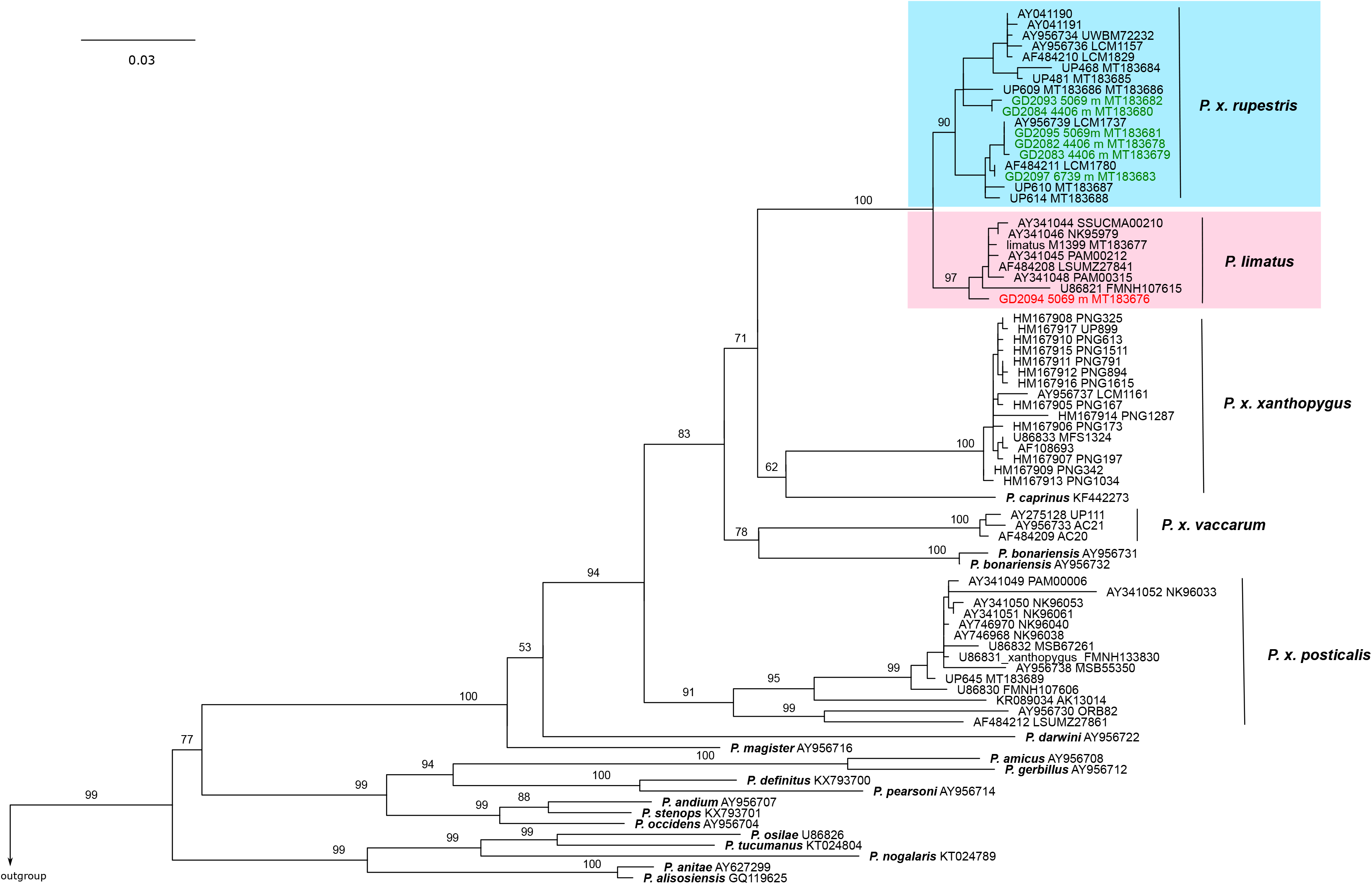
Phylogeny showing placement of high-altitude *Phyllotis* specimens from Volcán Llullaillaco. Majority rule consensus tree obtained in the Bayesian analysis of 76 *cytochrome-b* gene sequences from specimens of 18 species of *Phyllotis.* Numbers denote ultrafast Bootstrap support values for the adjacent nodes; only values for species clades and relationships among them are shown. Terminal labels are based on GenBank accession number and, when available, museum catalog number. Haplotypes of specimens collected at Volcán Llullaillaco are labeled in color.

In light of published suppositions about the altitudinal range limits of alpine mammals, our video footage and capture records are consistent with the view that “…the general dogma… about adaptation to the chronic hypoxia of high altitude has been ignored by the organisms proven most fit for life in such environments”^10^. Our capture of *P. x. rupestris* on the summit of Llullaillaco suggests that we may have generally underestimated the altitudinal range limits and physiological tolerances of small mammals simply because the world’s highest summits remain relatively unexplored by biologists. For logistical reasons, the elevational limits of most zoological collecting trips are largely dictated by road access. Consequently, the upper range limits of many vertebrate taxa are not precisely demarcated and putative altitudinal records for many taxa exist as unverified sightings or reports in mountaineering expedition accounts rather than as voucher specimens in museum collections.

Our discovery of the Llullaillaco summit mouse prompts many new evolutionary and ecological questions. Given the exceptionally broad altitudinal range of *Phyllotis xanthopygus* (from sea level to 6739 m), have mice from the upper reaches of Llullaillaco evolved genetically based adaptations to hypoxia that distinguish them from lowland conspecifics? To what extent is the ability to tolerate such a broad range of environmental conditions attributable to acclimatization (physiological plasticity)? Given that mice inhabiting the upper reaches of Llullaillaco are living >2000 m above the upper limits of green plants, what are they eating? Such questions can be answered by future mountaineering expeditions in the Humboldtian tradition that combine high-altitude exploration and scientific discovery.

## Materials and Methods

### Live-trapping and specimen preparation

We captured mice using Sherman live traps; the sole exception was the specimen from the Llullaillaco summit, which was captured by hand. We sacrificed mice in the field, prepared them as museum specimens, and preserved tissue samples as sources of DNA and RNA. Specimens are housed at the Colección de Mamíferos of the Universidad Austral de Chile, Valdivia, Chile. We identified all specimens to the species level based on cranial and post-cranial characters^20^, and all such identifications were subsequently confirmed with molecular sequence data. Additional tissue samples from Argentinean specimens were obtained as loans from the collection of the Centro Nacional Patagónico, Puerto Madryn, Argentina. Additional tissue samples from Argentinean and Peruvian specimens were obtained as loans from the collections of Centro Nacional Patagónico, Puerto Madryn, Argentina, and Louisiana State University Museum of Natural Science, Baton Rouge, USA.

All mice were collected in accordance with permissions to JFS from the following Chilean government agencies: Servicio Agrícola y Ganadero (SAG, Resolución extenta #209/2020), Corporación Nacional Forestal (CONAF, Autorización #’s 171219 and 1501221), and Dirección Nacional de Fronteras y Límites del Estado (DIFROL, Autorización de Expedición Cientifica #68). All mice were live-trapped and handled in accordance with protocols approved by the Institutional Animal Care and Use Committee (IACUC) at the University of Nebraska (project ID: 1919) and the American Society of Mammalogists^21^. Argentinean samples were exported with permission from the Dirección de Fauna y Flora Silvestres (export permit #3938/03).

### DNA sequencing

We sequenced the first 801 bp of the mitochondrial gene *Cytochrome-b* (*cytb*) from seven specimens of *Phyllotis* collected from three sites on Volcán Llullaillaco and 459-1144 bp of *cytb* from additional *Phyllotis* specimens collected in adjacent regions of Chile and Argentina. Sequences were generated following the protocols outlined by Steppan et al.^20^ and Teta et al.^22^, and automated DNA sequencing of the newly collected Llullaillaco specimens was conducted by AustralOmics (Valdivia, Chile). All new DNA sequences were deposited at GenBank (MT183676-MT183689).

### Taxon and character sampling for the phylogenetic analysis

We integrated newly generated sequences into a dataset containing GenBank sequences from *P. xanthopygus* as well as *P. bonariensi*s, *P. caprinus*, and *P. limatus* (three species that are nested within *P. xanthopygus* in *cytb* phylogenies)^19,23^. We excluded redundant sequences leaving one representative of each haplotype class. We also included in the matrix one representative sequence from every species of *Phyllotis* available in GenBank. The analyzed matrix consists of 76 sequences from 18 different species of *Phyllotis.* As outgroup taxa, we used *cytb* sequences from *Auslicomys pictus* and *Loxodontomys micropus*, species that belong to genera that are closely related to *Phyllotis*.

### Phylogenetic analysis

Sequences were aligned using Clustal W^24^, as implemented in Mega 7^25^. The best-fit model of nucleotide substitution (HKY+I+G) was determined based on the Bayesian Information Criterion (BIC) using jModeltest2^26^. We estimated the Maximum Likelihood *cytb* phylogeny using IQ-TREE^27^, as implemented in the IQ-TREE web server^28^, specifying the selected model of molecular evolution, with perturbation strength set to 0.5, and the number of unsuccessful iterations set to 100. Branch support was estimated via 1000 replicates of ultrafast Bootstrap^29^.

## Data availability

Data supporting the findings of this work are available within the paper and the Supplementary Information files. All sequence data have been deposited in GenBank and all voucher specimens have been cataloged in the Colección de Mamíferos of the Universidad Austral de Chile, Valdivia, Chile.

## Acknowledgments

This work was funded by grants to JFS from the National Institutes of Health (HL087216), National Science Foundation (OIA-1736249), and National Geographic Society (NGS-68495R-20), a grant to GD from the Fondo Nacional de Desarrollo Científico y Tecnológico (Fondecyt 1180366), and a grant to SJS from the National Science Foundation (DEB-0108422). We thank Ulyses Pardiñas, Mark Hafner, and Donna Dittmann for tissue loans from Argentinean *Phyllotis* specimens, Cristina Dorador for logistical support and guidance, and Mario Perez-Mamani and Juan Carlos Briceño for assistance and companionship in the field.

## Author contribution

Conceptualization, J.F.S., M.Q.-C., G.D.; Field work, J.F.S., M.Q.-C., G.D.; Investigation, J.F.S., M.Q.-C., J.C.O., G.D.; Resources, J.F.S., M.Q.-C., J.C.O., T.B., M.F., S.J.S., G.D; Writing – Original Draft, J.F.S., M.Q.-C., G.D; Writing – Review & Editing, J.F.S., M.Q.-C., J.C.O., T.B., M.F., S.J.S., G.D; Funding acquisition, J.F.S., G.D., S.J.S.

## Competing interests

The authors declare no competing interests.

Correspondence and requests for materials should be addressed to J.F.S. or G.D.

