## Supplemental Table 1 for "Discovery of the world’s highest-dwelling mammal"

Table S1. Trapping localities in the altiplano and Puna de Atacama, Regiόn de Antofagasta, northern Chile.

|  | Locality | Elevation | Coordinates |
| --- | --- | --- | --- |
| 1. | Chile, Antofagasta, San Pedro de Atacama, Ruta B-245 km 20 | 3240 m | 22º46.686’S, 68º04.619’W |
| 2. | Chile, Antofagasta, San Pedro de Atacama, Ruta 27 CH km 33 | 4099 m | 22º55.543’S, 67º52.383’W |
| 3. | Chile, Antofagasta, San Pedro de Atacama, Ruta 27 CH km 45.5 | 4750 m | 22º55.202’S, 67º46.042’W |
| 4. | Chile, Antofagasta, San Pedro de Atacama, Ruta B-241 km 117 | 2370 m | 23º01.095’S, 68º08.175’W |
| 5. | Chile, Antofagasta, Antofagasta, Parque Nacional Llullaillaco, 300 m al norte del Refugio Aguadas de Zorritas | 4140 m | 24°37.160’S, 68°35.325’W |
| 6. | Chile, Antofagasta, Antofagasta, Parque Nacional Llullaillaco, Refugio Aguadas de Zorritas | 4150 m | 24°37.293'S, 68°35.225'W |
| 7. | Chile, Antofagasta, Antofagasta, Parque Nacional Llullaillaco, 3 km al suroeste del Refugio Aguadas de Zorritas | 4360 m | 24°37.736’S, 68°33.580’W |
| 8. | Chile, Antofagasta, Antofagasta, Parque Nacional Llullaillaco, Volcán Llullaillaco, Campamento Base – Ruta Normal | 4620 m | 24°40.535’S, 68°34.843’W |
| 9. | Chile, Antofagasta, Antofagasta, Parque Nacional Llullaillaco, Volcán Llullaillaco, Campamento Base – Ruta Sur | 5070 m | 24°43.811’S, 68°34.676’W |
| 10. | Chile, Antofagasta, Antofagasta, Parque Nacional Llullaillaco, Volcán Llullaillaco, Campamento Alto – Ruta Sur | 5850 m | 24°43.288’S, 68°33.563’W |
| 11. | Chile, Antofagasta, Antofagasta, Parque Nacional Llullaillaco, cumbre del Volcán Llullaillaco | 6739 m | 24°43.235’S, 68°32.208’W |
